# Data aggregation strategies for a P300 speller: decoding models, epoch averaging, cross-subject ensembles, and multi-channel models

**DOI:** 10.64898/2026.06.17.732982

**Authors:** Leonid Sidorov, Anna Makarova, Archil Maysuradze, Mikhail Lebedev

## Abstract

Accurate detection of P300 event-related potentials from electroencephalography (EEG) remains challenging for small numbers of trials due to low signal-to-noise ratios and substantial inter-subject variability. This study presents a systematic comparison of data aggregation strategies for improving P300 classification, evaluated on a 10-subject dataset using two convolutional neural network architectures (EEGNet and BaseCNN) and a support vector machine (SVM). We compared: (1) subject-specific and pooled general models for single trials; (2) epoch averaging with 5 and 10 stimuli repetitions; (3) multi-channel models where subjects corresponded to different input channels; (4) cross-subject averaging; (5) mixed (uncontrolled) averaging; (6) a combined approach with *K* trials per subject across all participants; and (7) time-shifted channels from extended single-trial epochs. Decoding performance was quantified using the Information Transfer Rate (ITR), computed for binary classification accuracy. We found that single-trial ITR was unpractical (0.15–0.64 bits/trial), whereas controlled aggregation improved the performance. The combined cross-subject approach with *K* = 3 trials per participant (30 channels) achieves the highest ITR with multi-channel EEGNet: 0.95 bits/aggregated decision in the no-aperture recordings and 0.97 bits/aggregated decision on Aperture data, approaching the theoretical binary-classification limit for the aggregated decision. Controlled cross-subject averaging consistently outperformed random trial mixing, and multi-channel architectures out-performed simple averaging when inter-subject structure was preserved. These findings contribute to improving P300 decoding and implementing multi-subject brain-computer interfaces (BCIs).

## 1 Introduction

Event detection in electroencephalography (EEG) remains one of the most informative yet challenging problems in neural signal processing. EEG signals are noisy, and strongly variable across subjects and sessions, which complicates the construction of universal models for identifying cognitive events such as the P300 potential—a positive deflection in the event-related potential (ERP) that emerges around 300 ms after the presentation of a salient stimulus. The P300 component has long served as a benchmark paradigm for brain–computer interfaces (BCIs) and cognitive state monitoring because it reflects selective attention and decision-making processes [1, 2, 3]. The reliability and relative robustness of the P300 make it an attractive signal for such event-detection tasks. However, accurately detecting single-trial P300 responses amid EEG noise remains challenging, spurring extensive research into more effective classification algorithms.

Traditionally, P300 detection has relied on hand-crafted feature extraction and classical machine learning classifiers. In a typical pipeline, EEG signals are first preprocessed (filtered and segmented), then transformed to expose discriminative features of the P300. Spatial filtering methods like Common Spatial Patterns (CSP) or the xDAWN algorithm are often used to enhance the signal-to-noise ratio of ERPs by linearly combining channels [4]. Temporal features may be distilled via down-sampling or principal component analysis. These features are then fed into well-established classifiers such as Linear Discriminant Analysis (LDA) or support vector machines (SVM) for binary classification of “P300 vs. non-P300” (target vs. non-target) trials [5]. In particular, stepwise linear discriminant analysis (SWLDA) became a standard algorithm for P300 spellers, owing to its simplicity and strong performance [6]. Farwell and Donchin’s original BCI speller protocol included an SWLDA-based classifier, and subsequent studies found that LDA/SWLDA could outperform more complex methods like nonlinear SVMs on common P300 datasets. For example, Krusienski et al. compared several classifiers (Pearson correlation, LDA, SWLDA, linear and kernel SVM) on a P300 task and found that LDA and SWLDA achieved the highest accuracies [7]. These results underscored that a well-tuned linear classifier with appropriate feature engineering can be very effective for P300 detection. Classical methods also have the advantage of being data-efficient and interpretable. However, they depend heavily on human expertise to design features and may struggle with the non-stationary, complex dynamics of EEG signals.

In recent years, deep learning has become a powerful approach for EEG event detection, including P300 classification. Deep neural networks, and in particular convolutional neural networks (CNNs), can learn to extract and classify EEG features in an end-to-end manner, reducing the need for manual feature design [4]. Researchers have developed specialized CNN architectures for EEG: for example, Schirrmeister et al. (2017) introduced the DeepConvNet (and a related ShallowConvNet) for decoding motor and auditory EEG, achieving performance on par with the celebrated filter-bank CSP approach [8]. Lawhern et al. (2018) proposed EEGNet, a compact CNN utilizing depthwise separable convolutions to efficiently learn frequency–spatial features. EEGNet demonstrated high accuracy across multiple BCI paradigms—including P300 speller data—and importantly showed good generalization both within subjects and across subjects [9]. The success of networks like DeepConvNet and EEGNet indicated that deep models can automatically learn the pertinent spatiotemporal patterns (e.g. characteristic P300 waveforms over parietal electrodes) directly from raw data. Moreover, deep learning models have greater capacity to capture non-linear relationships and complex dependencies in EEG. This has led to state-of-the-art results in many EEG classification tasks. For instance, recent comparisons on a 75-subject dataset found that modern deep CNNs achieved high single-trial P300 detection accuracy, rivaling classic approaches [10]. However, deep methods typically require large training datasets and careful regularization; when only limited data are available per user (a common scenario), simpler methods like SWLDA can still match or exceed deep networks in within-subject performance [10]. This points to a trade-off: deep learning offers superior asymptotic performance with enough data, while classical methods may generalize better in data-sparse regimes.

A central challenge in EEG event detection is the inter-subject variability of brain signals. P300 responses can differ markedly between individuals in amplitude, latency, and scalp topography [11]. Even the same subject’s EEG varies across sessions due to electrode positioning, alertness, or other non-stationarities. This variability means that a model trained on one person’s data often performs poorly when applied to another person. In practice, BCI systems have traditionally handled this by using subject-specific models: each user undergoes a calibration session to record training data, and a personalized classifier is trained for that user. Subject-specific modeling maximizes performance because the classifier can fully adapt to an individual’s signal characteristics. The downside, however, is the lengthy calibration required for each new user and for each session, which is laborious and infeasible for real-time BCI deployment. There is strong motivation to develop subject-independent models that can generalize across users, eliminating or minimizing per-user training. A straightforward approach is to train a universal model on pooled data from many subjects and hope that it learns generalizable P300 features. In fact, deep learning models have shown promise in this regard: one study found that while a deep CNN did not outperform SWLDA in a within-subject scenario, the same CNN (EEG-Inception) significantly outperformed SWLDA in an across-subject training scenario [10]. This suggests that with enough diverse training data, a network can learn a representation of P300 that transfers to new individuals. However, building a truly subject-independent model is difficult because the network may instead learn the average pattern that dilutes important person-specific idiosyncrasies. Empirically, pooling data can sometimes even hurt performance on each individual, as the classifier must contend with greater between-subject variance. This limitation of universal models has driven researchers to explore intermediate solutions like domain adaptation and transfer learning to achieve better cross-subject generalization.

To mitigate inter-subject discrepancies, a variety of transfer learning (TL) and domain adaptation techniques have been proposed for EEG and P300-based BCIs. The key idea is to leverage knowledge from a set of source subjects to improve classification on a target subject, without requiring extensive target data. One class of approaches operates by aligning the feature distributions between subjects. For example, algorithms based on Riemannian geometry compute covariance matrices of EEG trials and then apply transformations to center these covariance features to a common reference, making different subjects more comparable [12]. Another prominent strategy is supervised fine-tuning of deep networks: one first trains a deep model on a large multi-subject dataset, then adapts it to a new subject by continuing training on a small amount of that subject’s data [13]. Beyond fine-tuning, there are also unsupervised domain adaptation methods that don’t require labeled target data: for example, extracting features in an unsupervised manner or using adversarial training to make the model’s features indistinguishable between source and target subjects. These advanced methods are actively being studied [14, 15, 16, 17, 18, 19], driven by the ultimate goal of zero- or minimal-calibration BCIs. In summary, cross-subject learning techniques now form a rich subfield, encompassing data alignment, augmentation, and transfer learning—all aiming to shrink the gap between individualized and universal modeling.

Ensemble models offer another promising approach to improve generalization in P300 detection. Ensemble learning combines multiple classifiers such that their individual errors can be averaged out, yielding a more robust overall prediction. A classic use of ensembles is to train several “weak” learners on different subsets or partitions of the data and then vote or average their outputs. This technique has been applied to P300 BCIs to boost performance: for example, Onishi and colleagues showed that an ensemble of LDA classifiers, each trained on a different portion of the ERP data, outperformed a single LDA, especially when an improved partitioning method was used to maximize data usage [6]. In fact, the winning entry of BCI Competition III (dataset II) was an ensemble of support vector machines that reduced EEG variability by averaging outputs from multiple SVM models [20]. In recent work, ensembles have also been explored to tackle subject variability. Mussabayeva et al. (2021) trained several classifiers—including LDA, SVM, kNN, and CNN—on data from a group of healthy subjects, and then used an ensemble voting strategy to classify signals from paralyzed patients without any retraining [11]. This subject-independent ensemble achieved over 90% spelling accuracy on the patients, about 5% higher than the best single classifier’s accuracy, underscoring that an aggregated model can be more stable against variations than any single model alone.

A complementary strategy to ensemble methods is signal-level aggregation through epoch averaging. In the classical ERP paradigm, multiple presentations of the same stimulus are averaged together before classification, exploiting the fact that the event-locked P300 component adds coherently while uncorrelated noise cancels out [21]. This approach is standard in P300 spellers, where several highlighting rounds are performed before making a character decision. The number of repetitions controls a trade-off between accuracy and communication speed: more repetitions improve signal quality but reduce the information throughput of the BCI. The Information Transfer Rate (ITR), introduced by Wolpaw et al. [22], captures this trade-off by quantifying the amount of information conveyed per trial in bits. While epoch averaging is well established, systematic investigations comparing different aggregation strategies—within-subject repetition averaging, cross-subject signal combination, and hybrid approaches—using modern deep learning architectures are scarce.

Despite the progress in each of the areas above, there is a need for testing different data aggregation strategies for P300 event detection. Prior studies tended to introduce a new method and evaluate it in isolation or against a few baselines, often under differing conditions, which makes it hard to discern the practical trade-offs. To address this gap, the present study conducts a controlled, head-to-head comparison of aggregation strategies for P300 detection: (1) subject-specific and general models on single trials, (2) multi-channel models treating multiple subjects as input channels, (3) epoch averaging with multiple stimulus repetitions per subject, (4) cross-subject averaging where each channel represents a different participant, (5) mixed averaging from an uncontrolled pool of trials, (6) a combined approach with *K* trials per subject from all participants, and (7) time-shifted channels constructed from overlapping windows within a single extended epoch. We evaluate all approaches on the same dataset using consistent preprocessing and three classifiers (BaseCNN, EEGNet, and SVM), with ITR as the primary metric. As detailed in Section 4, we systematically compare these strategies to determine which approach best balances performance, generalizability, and deployment assumptions. The findings, discussed in Section 5 and summarized in Section 6, provide practical guidelines for interpreting and choosing aggregation approaches under specific use-case requirements.

## 2 Dataset

We used a dataset comprising EEG recordings collected from ten subjects under identical conditions. The primary focus was the P300 potential, recorded under a classical P300 speller paradigm where the participant’s field of view was unrestricted. Within this framework, epochs during which the participant fixated attention on a highlighted symbol are considered targets. The data collection procedure is described in detail in [23]. The data collection interface is shown in Fig. 1.

**Figure 1:**
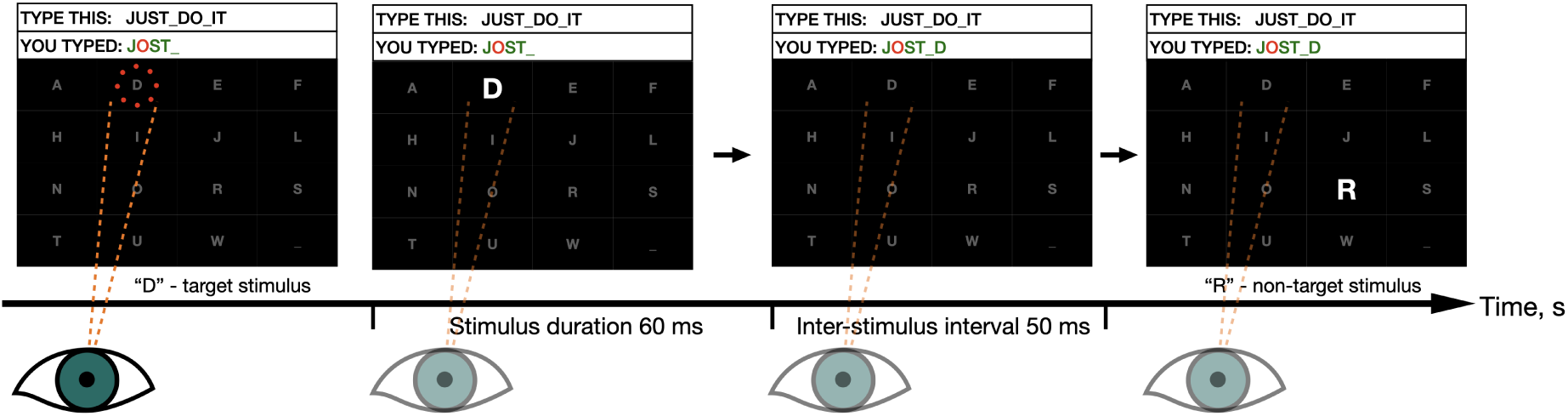
The P300 speller interface. Sixteen symbols are arranged in a 4 × 4 matrix, with the target phrase (“TYPE THIS”) and the currently decoded output (“YOU TYPED”) shown above the grid. The participant fixates gaze on the intended symbol (dashed lines and eye icon) while symbols are flashed one at a time: a flash of the attended symbol (“D”) is a target stimulus eliciting a P300 response, whereas a flash of any other symbol (“R”) is a non-target stimulus. Each stimulus lasts 60 ms followed by a 50 ms inter-stimulus interval (110 ms stimulus-onset asynchrony). The arrow indicates the progression of stimulus presentations over time.

The study involved 10 healthy subjects who had not previously been trained in the P300 BCI paradigm, all male, aged 19, right-handed, with the right dominant eye determined by the Miles test, with visual acuity of 1.04 0.18 (mean standard deviation) and 1.09 ± 0.18 for the left and right eye, respectively. Each participant was assigned a unique identifier from S0201 to S2001. The sampling rate was 250 Hz. Data were recorded from three primary EEG channels: P4, Pz, and P3. The recording was performed according to the international “10–20” system. The left earlobe was used as the reference channel, and the right earlobe as ground. The EEG was filtered with a 4th-order zero-phase Butterworth bandpass filter in the 1.0–15.0 Hz band. In this P300 BCI implementation, 16 symbols were displayed on the computer screen in a 4×4 matrix. Symbols were randomly highlighted one at a time. When highlighted, the symbol size doubled and its brightness increased by 80%. Stimuli (target and non-target) were presented every 110 ms (i.e., at a rate of 9.1 Hz). Stimulus duration was 60 ms. Participants used this P300 BCI to type a ten-character phrase (“JUST DO IT”). In the original study [23], classification was performed using an SVM classifier.

For the analyses, we used preprocessed data that had been segmented into fixed-length EEG epochs (250 samples, corresponding to 1 s) with corresponding class labels, where 1 indicates the presence of a target stimulus and hence a P300 response, and 0 indicates its absence. The recordings contain only the Pz channel, since it carries the primary signal information. Each epoch was standardized by centering and normalizing to unit variance.

In addition to the no-aperture (unrestricted visual field) recordings, the same experimental protocol was also performed under an *aperture condition* that restricted the subject’s visual field to the central area [23]. In this condition, participants wore a binocular aperture headset with a 5 mm opening placed 12 cm from the eyes, yielding an effective field of view of approximately 3.2–5.7^◦^ (depending on pupil diameter). With this restriction, subjects could see only one symbol at a time, which reduces interference from peripheral non-target flashes while keeping the same stimulus timing and EEG acquisition settings. In the remainder of the paper, we refer to this subset as *Aperture data* and use it as an additional replication condition.

## 3 Methods

### 3.1 Class Balancing: Borderline-SMOTE

The dataset exhibits significant class imbalance (1:16 ratio of target to non-target stimuli), necessitating the application of balancing techniques. We used the Borderline-SMOTE algorithm [24], an extension of SMOTE [25] that focuses synthetic minority sample generation on instances located near the decision boundary.

Borderline-SMOTE proceeds as follows: (1) for each minority sample, find its *m* nearest neighbors from the entire training set and count how many belong to the majority class; (2) categorize the minority sample as *Safe* (few majority neighbors), *Danger* (majority among the *m* neighbors exceeds *m/*2), or *Noise* (all neighbors are majority); (3) construct a DANGER set comprising only the Danger minority samples; (4) for each sample in the DANGER set, synthesize new minority examples by interpolating along line segments to its nearest minority neighbors. This border-focused strategy typically yields a greater increase in minority recall with a smaller rise in false positives compared to standard SMOTE. Given the extreme class imbalance (1:16), we used parameters *k* = 20, *m* = 40.

### 3.2 Evaluation Metric: Information Transfer Rate

The key evaluation metric used throughout this study is the Information Transfer Rate (ITR), defined by Wolpaw’s formula [22]:

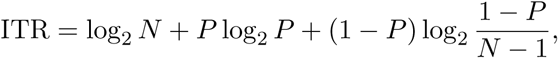

where *N* = 2 is the number of classes and *P* is the classification accuracy. ITR is measured in bits per evaluated decision unit and reflects the amount of information extracted by the classifier in information-theoretic units. For single-trial models, this unit is one stimulus presentation. For aggregation experiments, the unit is the aggregated sample formed from repeated trials, multiple subjects, or time-shifted windows, as specified in each subsection. The maximum ITR for binary classification is 1 bit per decision unit (at perfect accuracy). In the context of P300 BCI, ITR is a standard metric that allows comparison of decoding approaches while retaining the accuracy-throughput trade-off; operational communication rate in an online system additionally depends on stimulus timing, number of repetitions, and how trials are grouped before the final decision.

### 3.3 BaseCNN Architecture

The base model (Fig. 2) begins with a 1 × 1 pointwise convolution [26] applied across channels at each time step for inputs shaped [*C, T*]. This operation performs a learned linear projection that compresses the multivariate signal into a univariate time series while preserving temporal structure. Rather than averaging, the projection adaptively weights electrodes, emphasizing task-relevant sites, which reduces model complexity and overfitting, improves computational efficiency, and increases robustness to montage variability [27, 28, 29].

**Figure 2:**
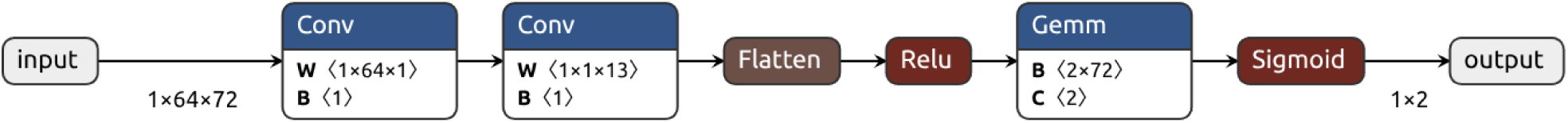
Base neural network architecture (BaseCNN).

A subsequent one-dimensional convolution along time models temporal dependencies; ReLU nonlinearities follow each convolution, and a final fully connected layer with sigmoid activation outputs class probabilities.

### 3.4 EEGNet Architecture

EEGNet [9] is a compact CNN for EEG decoding that embeds signal-processing priors to achieve data-efficient spatiotemporal feature learning under low signal-to-noise and small-sample conditions. Its architecture mirrors classical EEG analysis within a differentiable framework: (1) an initial temporal convolution per channel functions as a learnable bandpass filter to capture task-relevant rhythms; (2) a subsequent depthwise spatial convolution operates across electrodes to learn discriminative spatial filters analogous to common spatial patterns, with a depth multiplier enabling multiple spatial projections per frequency component; (3) a separable convolution stage (depthwise temporal followed by pointwise 1 × 1) performs nonlinear feature mixing with minimal parameter growth. Training is stabilized through batch normalization, ELU activations, average pooling, and dropout, while the omission of large fully connected layers constrains model capacity (Fig. 3).

**Figure 3:**
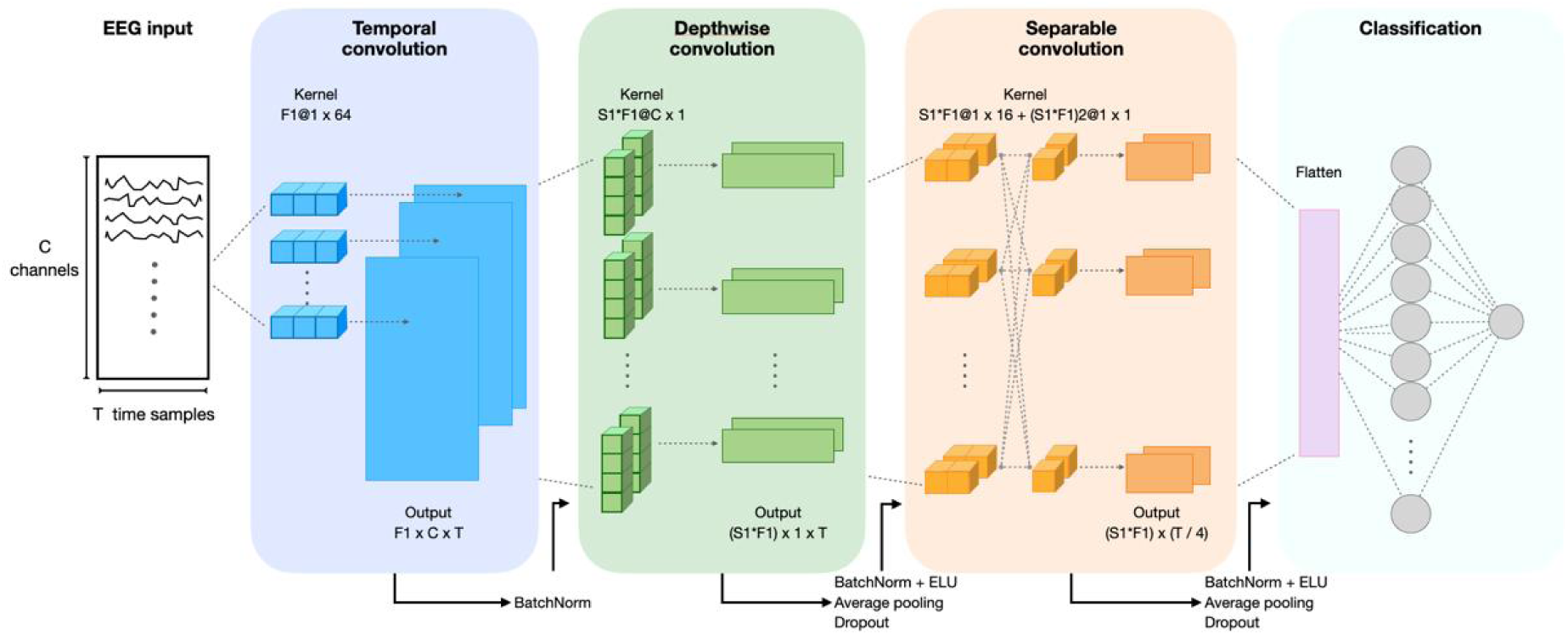
EEGNet architecture. An EEG input of *C* channels and *T* time samples passes through a temporal convolution (*F*1 × *C × T*), a depthwise spatial convolution ((*S*1 × *F*1) × 1 ×*T*), and a separable convolution ((*S*1 *F*1) × (*T/*4)), followed by flattening and a fully connected layer for binary classification. The depth multiplier is denoted *S*1 in the figure (equivalently *D* in the text). See Section 3.4 for details.

This design confers several practical benefits: parameter efficiency mitigates overfitting and supports reliable learning from limited datasets; end-to-end optimization on raw or minimally pre-processed signals reduces reliance on handcrafted features; a single backbone applies across major BCI paradigms with minimal retuning; and the learned temporal and spatial filters are amenable to neurophysiological interpretation [9].

### 3.5 Support Vector Machine

An SVM with a radial basis function (RBF) kernel was used as an additional comparison method. The SVM was applied to the same raw EEG data without additional preprocessing. For the multi-channel and averaging experiments, the SVM operated on the averaged (single-channel) representation, since it does not natively support multi-channel temporal inputs. The model was trained using the same train/validation splits as the neural network models.

## 4 Tests

The dataset for each subject was standardized to the minimum number of epochs across all subjects to ensure consistent input dimensions. We define an epoch as a complete cycle of screen highlighting during EEG recording, comprising one positive signal and sixteen negative signals. Train–validation splitting was performed before oversampling and aggregation. Borderline-SMOTE was applied only to the training data, and aggregated training and validation tensors were then constructed separately within their respective splits. This protocol prevents synthetic samples or source epochs from crossing the train–validation boundary.

Several tests below use class-conditioned aggregation: trials with the same known label are grouped to estimate the information gain produced by combining independent realizations of the same class. This setup is useful as a controlled offline comparison of aggregation strategies, but it is not itself a direct online BCI procedure because positive and negative class labels are unknown before classification. The practical online analogue in a P300 speller is to group responses by repeated presentations of the same candidate stimulus or symbol before making the final decision. Figure 4 summarizes the complete workflow and clarifies how the aggregation strategies are connected to the downstream model families. This visualization helps distinguish settings where aggregation is explicitly imposed (SC averaging) from settings where aggregation is learned end-to-end by multi-channel neural architectures (MC). Accuracy is reported together with ITR in the main tables. F1-score was monitored during hyperparameter tuning as a secondary imbalance-sensitive diagnostic, but ITR was retained as the primary metric because it directly reflects the BCI decision objective. The F1-score values followed the same qualitative trend as ITR in the repetition-averaging experiments, increasing with the number of aggregated stimulus repetitions, especially in the Aperture condition. The reported 95% confidence intervals are binomial proportion intervals for accuracy, computed from the number of correct validation decisions and the validation-set size.

**Figure 4:**
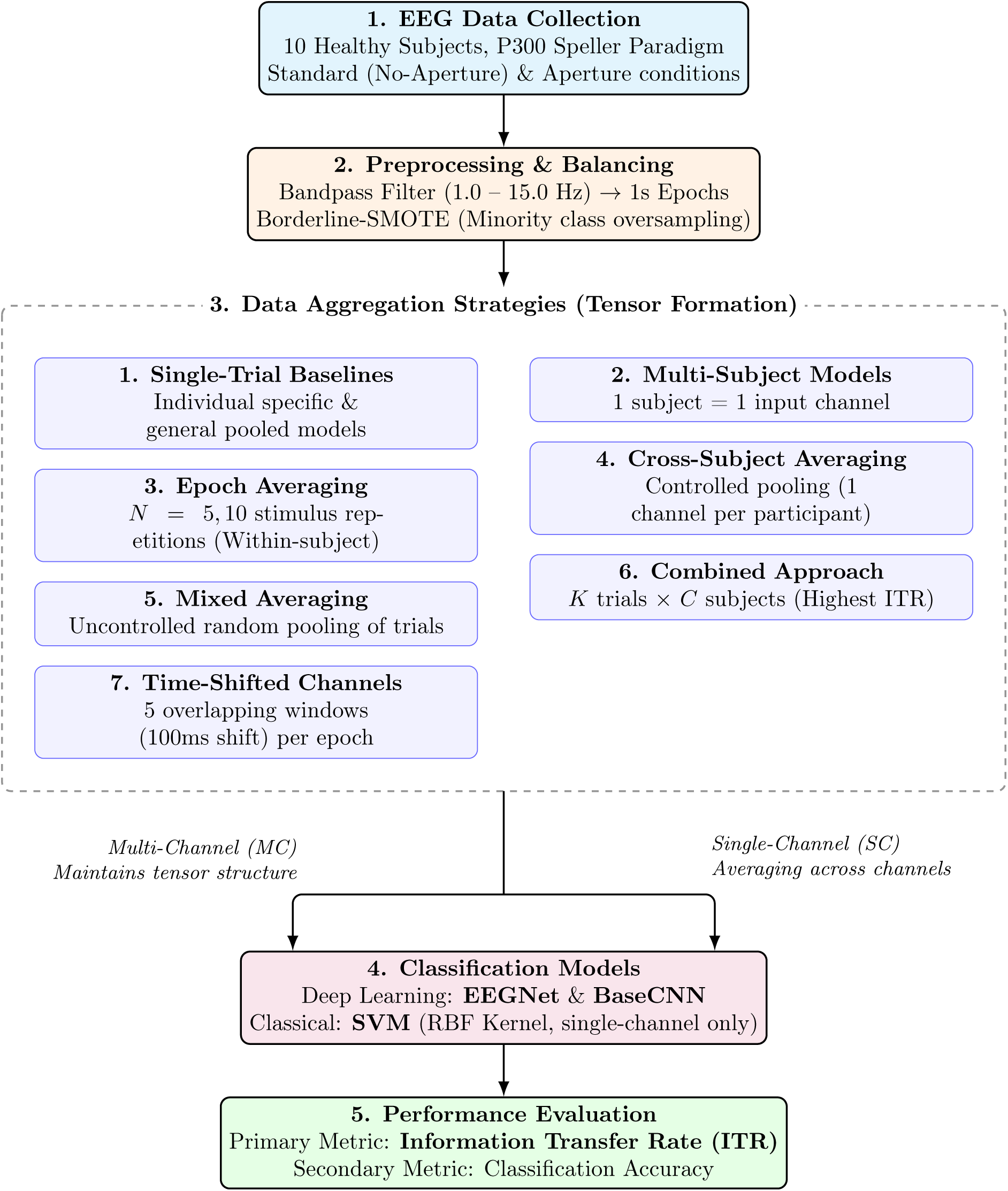
End-to-end experimental workflow and taxonomy of aggregation strategies evaluated in this study.

### 4.1 Subject-Specific Models

In the first experiment, individual models were trained and validated separately for each participant. This approach allows each model to adapt to subject-specific signal characteristics. Each input sample represents a univariate time series of length 250 (*x* ∈ R^250^). The task is binary classification of single stimulus presentations (target vs. non-target). Table 1 presents the results.

**Table 1:** Metrics of subject-specific P300 detection models. Acc. = accuracy, ITR = information transfer rate (bits/trial).

| Participant | EEGNet |  | BaseCNN |  | SVM |  |
| --- | --- | --- | --- | --- | --- | --- |
|  | Acc. | ITR | Acc. | ITR | Acc. | ITR |
| S0201 | 0.83 | 0.34 | 0.68 | 0.10 | 0.91 | 0.56 |
| S0601 | 0.80 | 0.28 | 0.77 | 0.22 | 0.93 | 0.63 |
| S0701 | 0.82 | 0.32 | 0.78 | 0.24 | 0.93 | 0.63 |
| S1201 | 0.84 | 0.37 | 0.69 | 0.11 | 0.92 | 0.60 |
| S1401 | 0.83 | 0.34 | 0.64 | 0.06 | 0.92 | 0.60 |
| S1601 | 0.87 | 0.44 | 0.77 | 0.22 | 0.92 | 0.60 |
| S1701 | 0.83 | 0.34 | 0.67 | 0.09 | 0.94 | 0.67 |
| S1801 | 0.82 | 0.32 | 0.67 | 0.09 | 0.93 | 0.63 |
| S1901 | 0.90 | 0.53 | 0.84 | 0.37 | 0.97 | 0.81 |
| S2001 | 0.79 | 0.26 | 0.64 | 0.06 | 0.93 | 0.63 |
| **Mean** | **0.83** | **0.35** | **0.72** | **0.15** | **0.93** | **0.64** |

Individual models showed high variability in decoding quality across participants, reflecting differences in the magnitude and stability of the P300 response. SVM showed the highest ITR among individual models (0.64 bits/trial) owing to its high accuracy (0.93). The mean ITR for EEGNet was 0.35 bits/trial, and for BaseCNN it was 0.15 bits/trial. For all participants, EEGNet outperformed BaseCNN in ITR, demonstrating the advantage of a specialized architecture for EEG tasks. The ITR for neural network models remained substantially below the maximum of 1 bit/trial, indicating that single-trial classification has limited practical utility. For practical applications, multiple repetitions of the target stimulus are necessary.

### 4.2 General (Pooled) Model

In the next experiment, a single generalized model was trained on the combined data of all users. The model classified single stimulus presentations, as in the previous analysis. Table 2 shows the results.

**Table 2:**
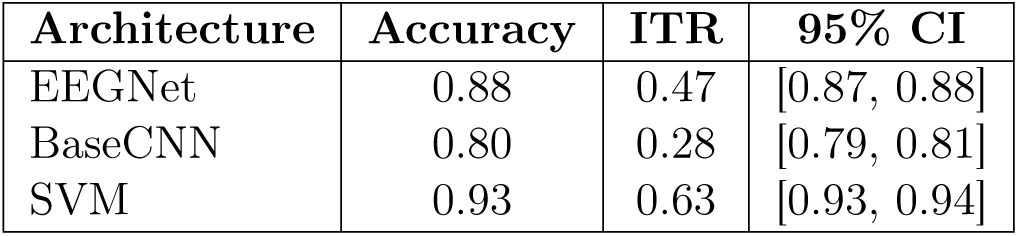
Performance of general models trained on pooled data from all subjects. CI denotes 95% confidence interval for accuracy.

SVM showed the highest ITR among generalized models (0.63 bits/trial, accuracy 0.93), outperforming EEGNet (ITR = 0.47) and BaseCNN (ITR = 0.28). However, the ITR of all generalized models remained below 0.65 bits/trial, confirming the need for stimulus repetitions for practical applications. Notably, the generalized SVM achieved ITR comparable to the mean of individual SVM models (0.64), suggesting SVM features are more robust to inter-subject variability than CNN features.

### 4.3 Multi-Subject Model

In the following experiment, signals from all users were combined into a single input tensor, where each user was represented by a separate channel. The model automatically learned optimal weights for each user, reflecting their relative contribution to P300 detection. This approach should be interpreted as a controlled multi-source aggregation setting: it is analogous to combining repeated observations, but the observations come from different participants rather than from the same user. Table 3 presents the results.

**Table 3:** Performance of multi-subject models where each participant serves as an input channel (10 channels total). ITR is reported in bits per evaluated decision unit.

| Architecture | Accuracy | ITR | 95% CI |
| --- | --- | --- | --- |
| EEGNet | 0.98 | 0.86 | [0.98, 0.99] |
| BaseCNN | 0.97 | 0.81 | [0.96, 0.97] |

The multi-channel EEGNet model achieved an ITR of 0.86 bits/decision unit (accuracy 0.983), a significant improvement compared to individual models (ITR = 0.35). This indicates that structured multi-source aggregation can substantially increase discriminability under controlled conditions. Importantly, averaging was not algorithmically prescribed; the neural network architectures learned to optimally combine the multi-channel input.

### 4.4 Learned Channel Weights

To interpret how the multi-channel models weight different subjects, we analyzed the learned spatial filters. For EEGNet, we extracted the effective channel weights by combin_√_in<u>g the</u> depthwise convolution weights with batch normalization scaling factors: *W*_eff_ = (*W^T^* ⊙ *γ/ σ*^2^ + *ɛ*)*^T^*, where *W* are the depthwise convolution weights, *γ* and *σ*^2^ are the batch normalization scale and variance parameters, respectively. For BaseCNN, we examined the pointwise convolution weights that project the multi-channel input to a univariate time series.

Figures 5 and 6 visualize the aggregated channel weights. Subjects S1901, S1601, and S1401 receive consistently high weights in both models, with S1901 achieving the highest weight in EEGNet (0.405). This aligns with the subject-specific results, where these subjects achieved the highest individual ITR values (S1901: 0.53 for EEGNet, 0.81 for SVM). Conversely, subjects S0701 and S2001 receive lower weights, consistent with their lower subject-specific performance.

**Figure 5:**
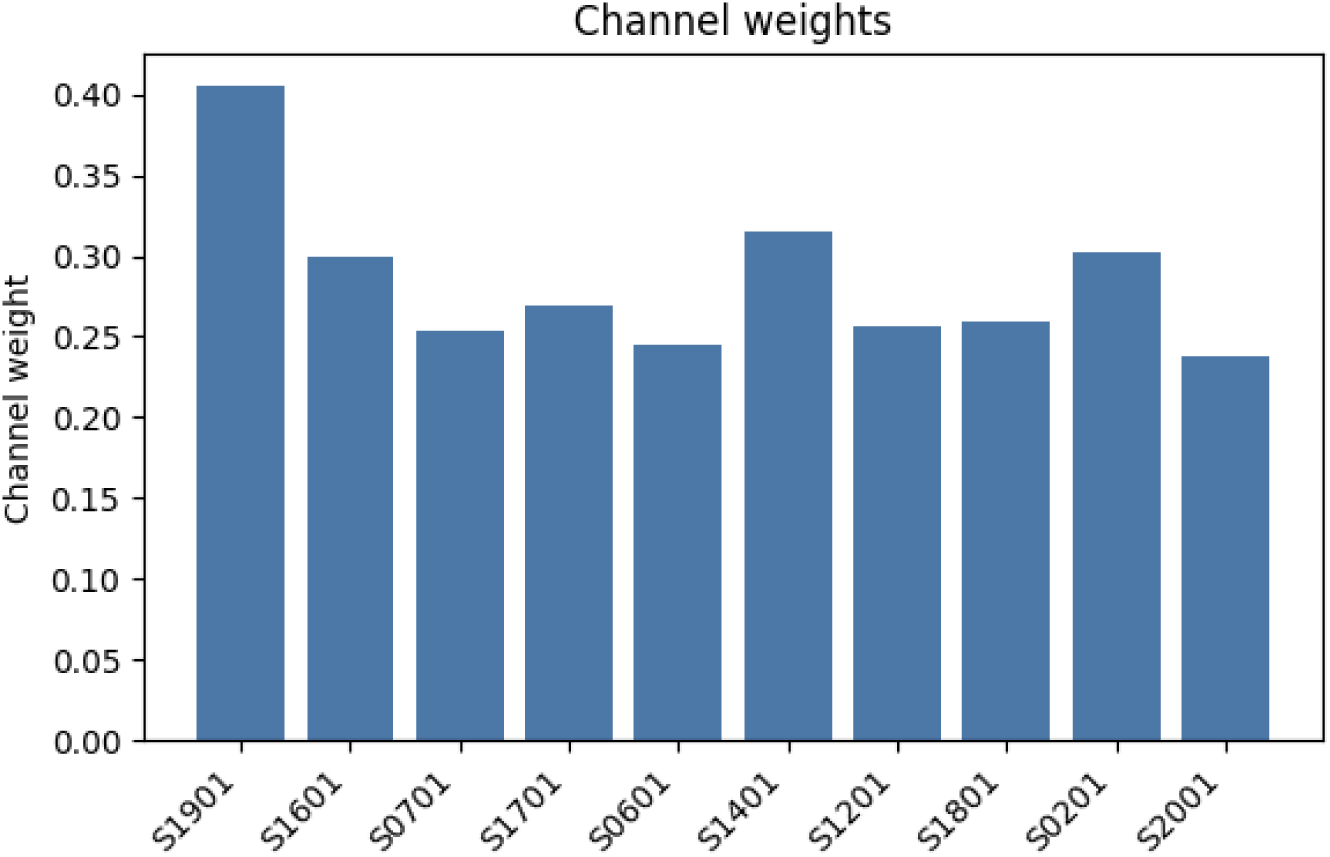
Aggregated channel weights learned by the multi-channel EEGNet model. Subjects S1901, S1401, and S1601 receive the highest weights, indicating their EEG patterns are particularly discriminative for P300 detection.

**Figure 6:**
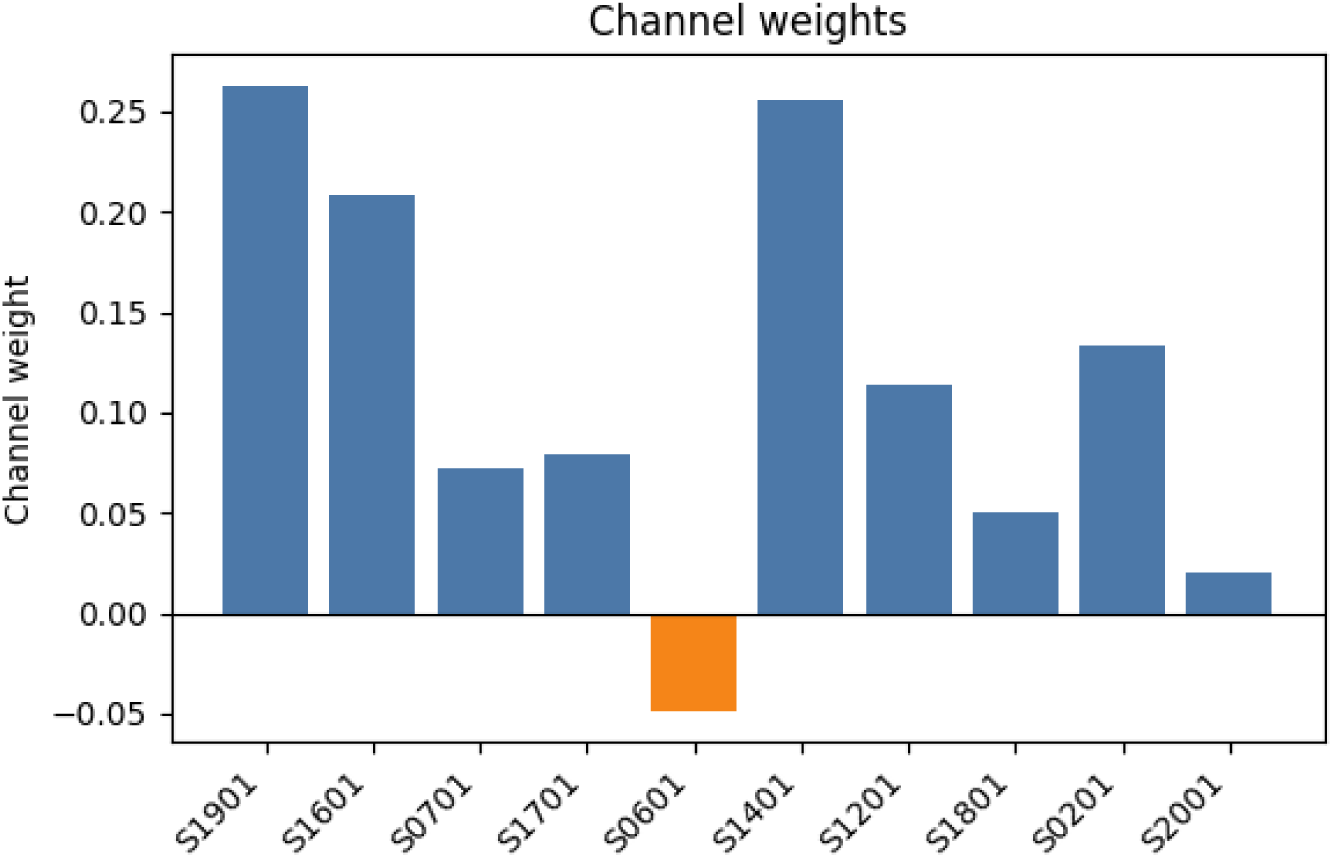
Channel weights learned by the multi-channel BaseCNN model. Subject S0601 receives a negative weight, suggesting its signal pattern may be inversely correlated with P300 responses in the learned representation.

BaseCNN learns a single linear combination with one subject (S0601) receiving a negative weight, indicating an inverse correlation with P300 responses. EEGNet’s aggregated weights are all positive, ranging from 0.24 to 0.405, suggesting a more uniform positive contribution with varying emphasis. Per-kernel analysis (Fig. 7) reveals that EEGNet learns a rich, multi-faceted representation where different kernels specialize in different aspects of the multi-subject P300 signal.

**Figure 7:**
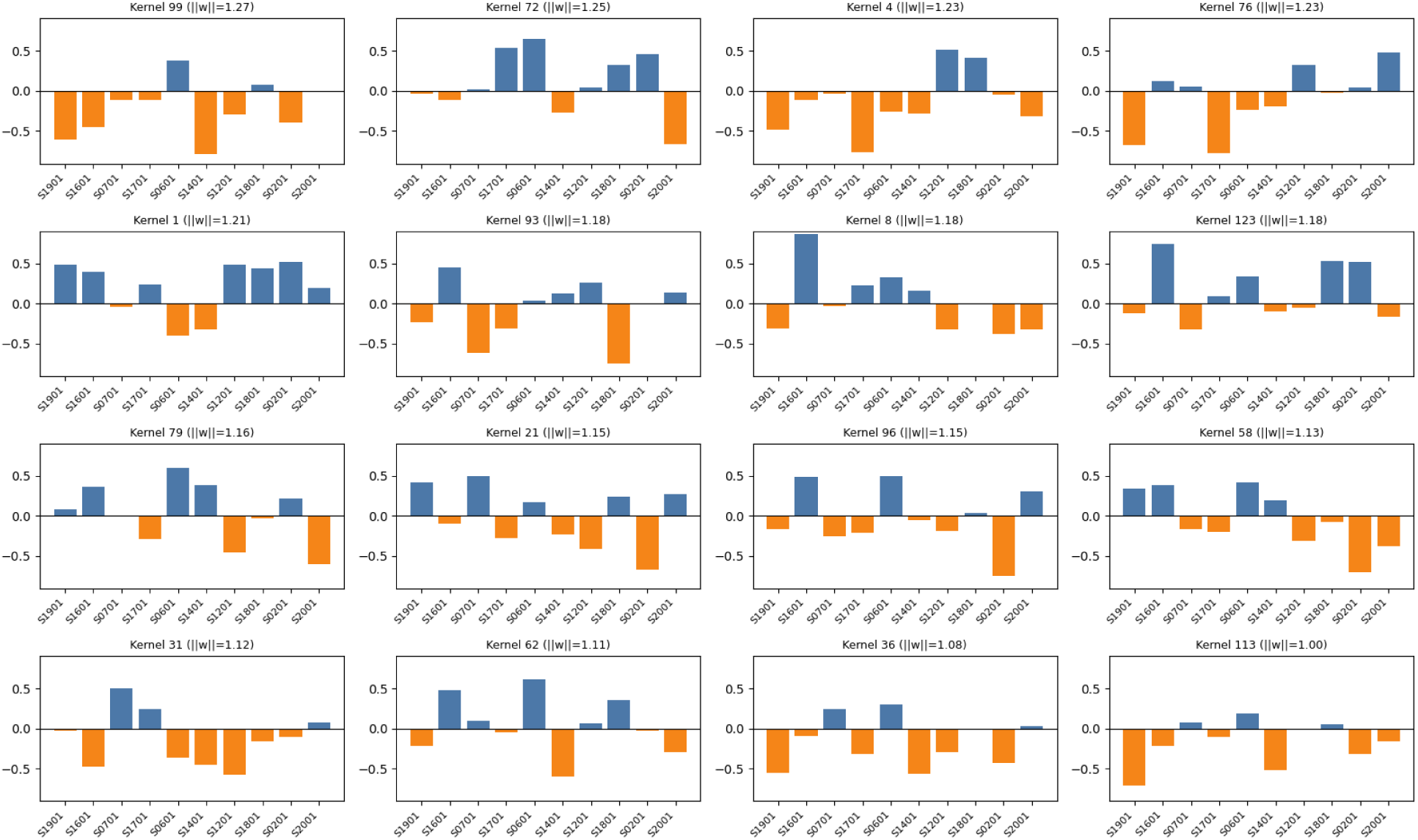
Channel weights for 16 representative kernels learned by the multi-channel EEGNet model. Each subplot shows the weights assigned by a single kernel to each subject. Kernels exhibit diverse weighting patterns, demonstrating that EEGNet learns multiple specialized spatial filters.

This adaptive weighting mechanism is a key advantage of the multi-channel approach: the model automatically identifies and emphasizes subjects whose P300 responses are most consistent and discriminative, effectively performing a learned weighted ensemble rather than uniform averaging.

### 4.5 Epoch Averaging: Multiple Stimulus Repetitions

In the next set of experiments, multiple class-conditioned stimulus presentations were used. To form the multi-channel input, several randomly selected stimulus presentations of the same class from the same subject were distributed across channels. This controlled offline construction estimates the gain from combining independent realizations of a target or non-target response. In an online P300 speller, the corresponding deployable operation would be averaging responses to repeated presentations of the same candidate stimulus before deciding whether it is the intended target. Experiments were conducted for three methods—BaseCNN, EEGNet, and SVM—using *N* = 5 and *N* = 10 channels (stimulus repetitions). For neural network architectures, two approaches were evaluated:

- **Multi-channel (MC):** the model received *N* channels as input, where each channel contained a separate stimulus presentation;
- **Single-channel (SC):** the same data were averaged across channels, and the model operated on the averaged representation.

For SVM, only the averaged (SC) representation was used.

Figure 8 illustrates the tensor construction for this within-subject setting. In the MC variant, each repetition is preserved as a separate input channel, allowing the model to learn non-uniform weights across repetitions. In the SC variant, repetitions are collapsed by arithmetic averaging before classification.

**Figure 8:**
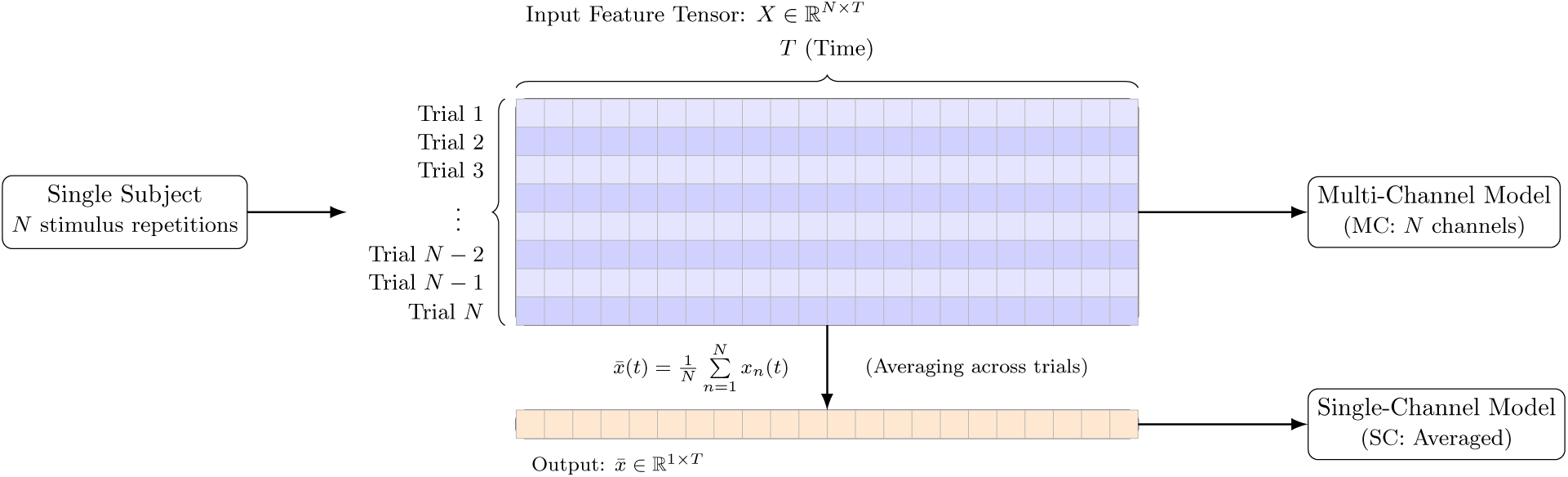
Within-subject epoch aggregation: MC preserves repetition-wise channels, while SC averages repetitions before classification.

The results are presented in Tables 4 and 5. Classification quality improved substantially when using multiple stimulus repetitions. When increasing the number of channels from 5 to 10, ITR increased by 6–37% depending on the model and approach. With 10 channels, the SC EEGNet achieved a mean ITR of 0.64 bits/aggregated decision, and SC BaseCNN achieved 0.58 bits/aggregated decision. The SC model with averaging showed comparable or higher ITR than the MC model, indicating that simple averaging is an effective aggregation method for within-subject repetitions.

**Table 4:** Multi-channel (MC) models with multiple stimulus repetitions per channel. ITR is reported in bits per aggregated decision.

| Participant | 5 channels |  |  |  | 10 channels |  |  |  |
| --- | --- | --- | --- | --- | --- | --- | --- | --- |
|  | BaseCNN |  | EEGNet |  | BaseCNN |  | EEGNet |  |
|  | Acc. | ITR | Acc. | ITR | Acc. | ITR | Acc. | ITR |
| S0201 | 0.90 | 0.52 | 0.92 | 0.60 | 0.93 | 0.63 | 0.92 | 0.60 |
| S0601 | 0.76 | 0.21 | 0.82 | 0.31 | 0.81 | 0.30 | 0.83 | 0.35 |
| S0701 | 0.76 | 0.20 | 0.85 | 0.38 | 0.82 | 0.32 | 0.83 | 0.34 |
| S1201 | 0.88 | 0.47 | 0.87 | 0.45 | 0.88 | 0.47 | 0.84 | 0.37 |
| S1401 | 0.90 | 0.52 | 0.93 | 0.62 | 0.91 | 0.57 | 0.89 | 0.50 |
| S1601 | 0.90 | 0.52 | 0.92 | 0.60 | 0.96 | 0.76 | 0.92 | 0.60 |
| S1701 | 0.80 | 0.27 | 0.84 | 0.37 | 0.87 | 0.45 | 0.90 | 0.53 |
| S1801 | 0.77 | 0.23 | 0.88 | 0.46 | 0.91 | 0.56 | 0.93 | 0.63 |
| S1901 | 0.96 | 0.74 | 0.98 | 0.83 | 0.98 | 0.86 | 0.98 | 0.86 |
| S2001 | 0.80 | 0.28 | 0.79 | 0.26 | 0.89 | 0.50 | 0.85 | 0.39 |
| **Mean** | **0.84** | **0.40** | **0.88** | **0.49** | **0.90** | **0.54** | **0.89** | **0.52** |

**Table 5:**
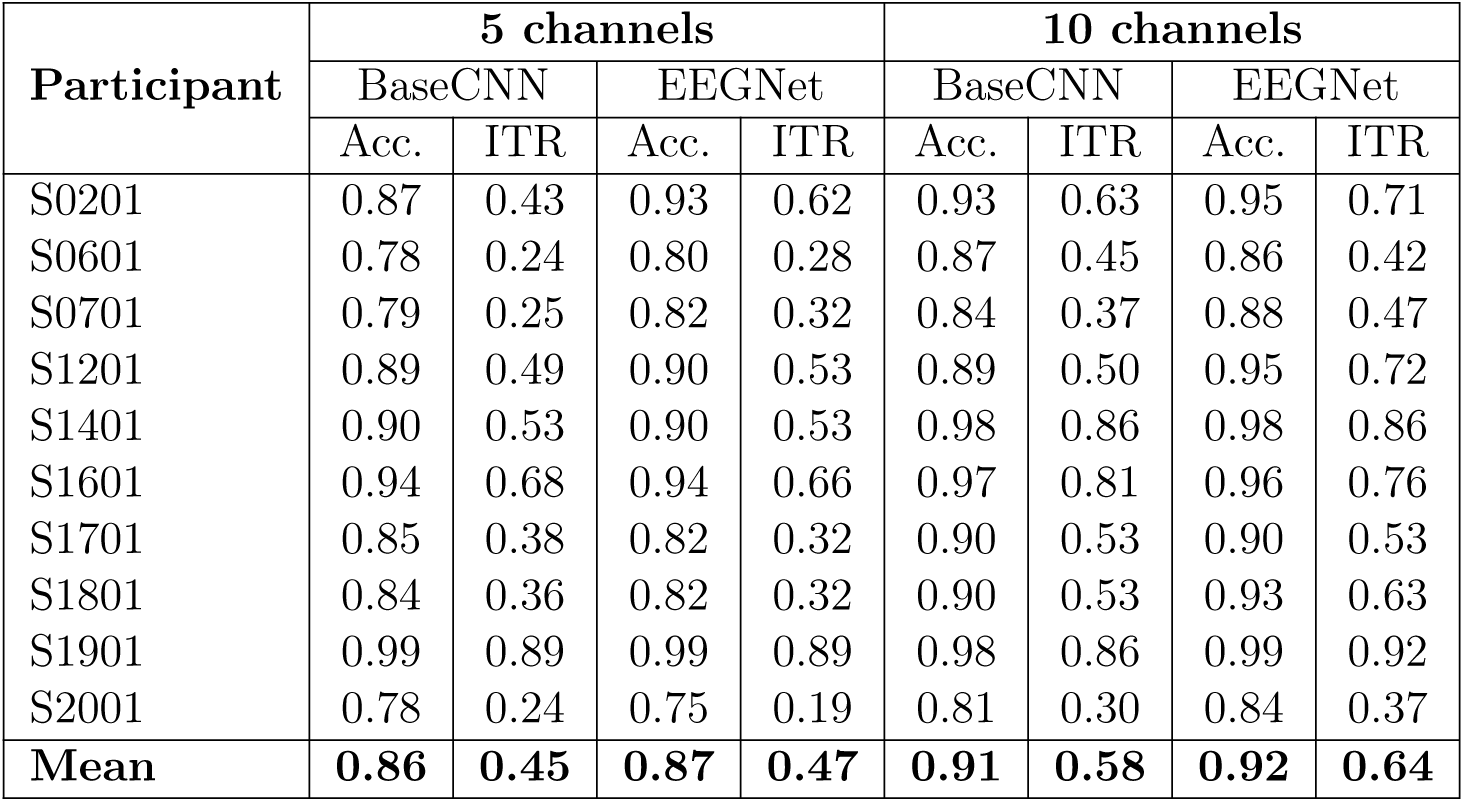
Single-channel (SC) models with averaging over 5 and 10 repetitions. ITR is reported in bits per aggregated decision.

| Participant | 5 channels |  |  |  | 10 channels |  |  |  |
| --- | --- | --- | --- | --- | --- | --- | --- | --- |
|  | BaseCNN |  | EEGNet |  | BaseCNN |  | EEGNet |  |
|  | Acc. | ITR | Acc. | ITR | Acc. | ITR | Acc. | ITR |
| S0201 | 0.87 | 0.43 | 0.93 | 0.62 | 0.93 | 0.63 | 0.95 | 0.71 |
| S0601 | 0.78 | 0.24 | 0.80 | 0.28 | 0.87 | 0.45 | 0.86 | 0.42 |
| S0701 | 0.79 | 0.25 | 0.82 | 0.32 | 0.84 | 0.37 | 0.88 | 0.47 |
| S1201 | 0.89 | 0.49 | 0.90 | 0.53 | 0.89 | 0.50 | 0.95 | 0.72 |
| S1401 | 0.90 | 0.53 | 0.90 | 0.53 | 0.98 | 0.86 | 0.98 | 0.86 |
| S1601 | 0.94 | 0.68 | 0.94 | 0.66 | 0.97 | 0.81 | 0.96 | 0.76 |
| S1701 | 0.85 | 0.38 | 0.82 | 0.32 | 0.90 | 0.53 | 0.90 | 0.53 |
| S1801 | 0.84 | 0.36 | 0.82 | 0.32 | 0.90 | 0.53 | 0.93 | 0.63 |
| S1901 | 0.99 | 0.89 | 0.99 | 0.89 | 0.98 | 0.86 | 0.99 | 0.92 |
| S2001 | 0.78 | 0.24 | 0.75 | 0.19 | 0.81 | 0.30 | 0.84 | 0.37 |
| **Mean** | **0.86** | **0.45** | **0.87** | **0.47** | **0.91** | **0.58** | **0.92** | **0.64** |

SVM with epoch averaging (Tables 6 and 7) demonstrated high effectiveness. Individual SVM models achieved a mean ITR of 0.57 bits/aggregated decision with 5 repetitions and 0.68 bits/aggregated decision with 10 repetitions. For four participants (S1201, S1401, S1601, S1901), ITR with 10 repetitions exceeded 0.76 bits/aggregated decision, and participant S1901 achieved perfect classification (ITR = 1.00). The general SVM model showed lower results (ITR = 0.37 and 0.54 for 5 and 10 channels), reflecting inter-subject EEG variability.

**Table 6:** SVM with epoch averaging (subject-specific models). ITR is reported in bits per aggregated decision.

| Participant | 5 channels |  | 10 channels |  |
| --- | --- | --- | --- | --- |
|  | Accuracy | ITR | Accuracy | ITR |
| S0201 | 0.931 | 0.64 | 0.940 | 0.67 |
| S0601 | 0.866 | 0.43 | 0.901 | 0.53 |
| S0701 | 0.842 | 0.37 | 0.860 | 0.42 |
| S1201 | 0.926 | 0.62 | 0.970 | 0.81 |
| S1401 | 0.936 | 0.66 | 0.960 | 0.76 |
| S1601 | 0.955 | 0.74 | 0.990 | 0.92 |
| S1701 | 0.886 | 0.49 | 0.901 | 0.53 |
| S1801 | 0.842 | 0.37 | 0.920 | 0.60 |
| S1901 | 0.995 | 0.96 | 1.000 | 1.00 |
| S2001 | 0.847 | 0.38 | 0.920 | 0.60 |
| **Mean** | **0.903** | **0.57** | **0.936** | **0.68** |

**Table 7:** SVM with epoch averaging (general model trained on pooled data). ITR is reported in bits per aggregated decision.

| Number of channels | Accuracy | ITR | 95% CI |
| --- | --- | --- | --- |
| 5 | 0.843 | 0.37 | [0.827, 0.858] |
| 10 | 0.903 | 0.54 | [0.885, 0.922] |

**Table 8:**
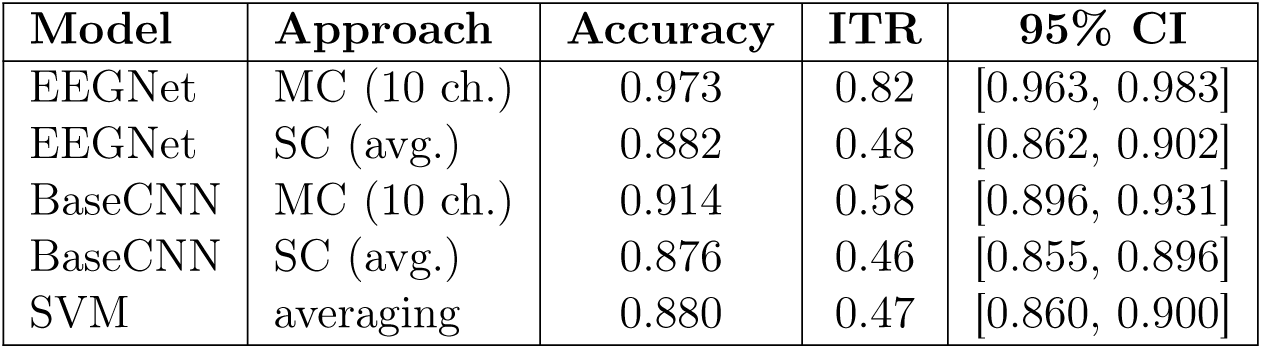
Cross-subject averaging (10 participants, each as a separate channel). ITR is reported in bits per aggregated decision.

### 4.6 Cross-Subject Averaging

In the following experiment, the input tensor was formed from data of all participants: each channel corresponded to one participant (10 channels in total). For each sample, stimulus presentations of the same class from different participants were combined. The model simultaneously processed signals from all 10 participants, allowing it to learn optimal weights for weighted averaging of individual responses. The number of samples was determined by the minimum number of trials per class across all participants. This experiment is best interpreted as controlled multi-source aggregation rather than a standard single-user BCI deployment scenario. For each architecture, both MC (10 channels) and SC (mean across participants) approaches were evaluated.

Multi-channel EEGNet achieved an ITR of 0.82 bits/aggregated decision, comparable to the multi-subject model from Table 3 (ITR = 0.86). The MC approach significantly outperformed SC: for EEGNet, the ITR difference was 0.34 bits/aggregated decision (0.82 vs. 0.48), indicating the model’s ability to extract additional information from inter-subject differences rather than averaging them out. SVM with averaging showed an ITR of 0.47 bits/aggregated decision, lower than its individual results with 10 repetitions (0.68), since each channel represented a different participant rather than a repetition of the same stimulus.

### 4.7 Mixed Averaging

A mixed averaging strategy was also investigated, in which all data from all participants were pooled, and groups of *N* trials (*N* = 5 and *N* = 10) were randomly formed from this pool. Unlike the cross-subject approach, a single group could contain multiple trials from one participant and none from another—subject assignment was not controlled. This models a situation where the homogeneity of data sources is not guaranteed.

Mixed averaging (Table 9) showed results substantially lower than the cross-subject approach: the best ITR was 0.51 bits/aggregated decision (BaseCNN SC, *N* = 10) versus 0.82 bits/aggregated decision for cross-subject EEGNet MC. This indicates that controlled combination of signals from different participants (one channel per subject) is more effective than random trial mixing. SC models with averaging generally performed comparably to MC models, consistent with the epoch averaging results.

**Table 9:** Mixed averaging (*N* random trials from the common pool). ITR is reported in bits per aggregated decision.

| Model | $N$ | Approach | Accuracy | ITR | 95% CI |
| --- | --- | --- | --- | --- | --- |
| EEGNet | 5 | MC | 0.771 | 0.22 | [0.752, 0.789] |
| EEGNet | 5 | SC | 0.830 | 0.34 | [0.814, 0.847] |
| BaseCNN | 5 | MC | 0.821 | 0.32 | [0.805, 0.838] |
| BaseCNN | 5 | SC | 0.802 | 0.28 | [0.784, 0.819] |
| SVM | 5 | averaging | 0.821 | 0.32 | [0.804, 0.838] |
| EEGNet | 10 | MC | 0.860 | 0.42 | [0.838, 0.881] |
| EEGNet | 10 | SC | 0.891 | 0.50 | [0.872, 0.911] |
| BaseCNN | 10 | MC | 0.885 | 0.48 | [0.865, 0.904] |
| BaseCNN | 10 | SC | 0.893 | 0.51 | [0.874, 0.912] |
| SVM | 10 | averaging | 0.889 | 0.50 | [0.870, 0.909] |

### 4.8 Cross-Subject Averaging with Multiple Trials

Finally, a combined approach was investigated that merged cross-subject averaging with multiple class-conditioned stimulus repetitions. From each participant, *K* trials (*K* = 2, 3) were taken, forming an input tensor of *N*_subjects_ × *K* channels (20 and 30 channels, respectively). Each sample was guaranteed to include representation from all participants, with each participant contributing *K* independent trials. Because this construction uses known class labels and multiple sources, it should be interpreted as an upper-bound controlled aggregation condition rather than a directly deployable online procedure.

Figure 9 shows this construction explicitly. Compared to simple cross-subject averaging (one channel per participant), this strategy enriches each subject-specific contribution with multiple independent realizations, thereby increasing the effective signal-to-noise ratio while preserving source identity. This dual structure explains why the MC models in Table 10 achieve the highest ITR values.

**Figure 9:**
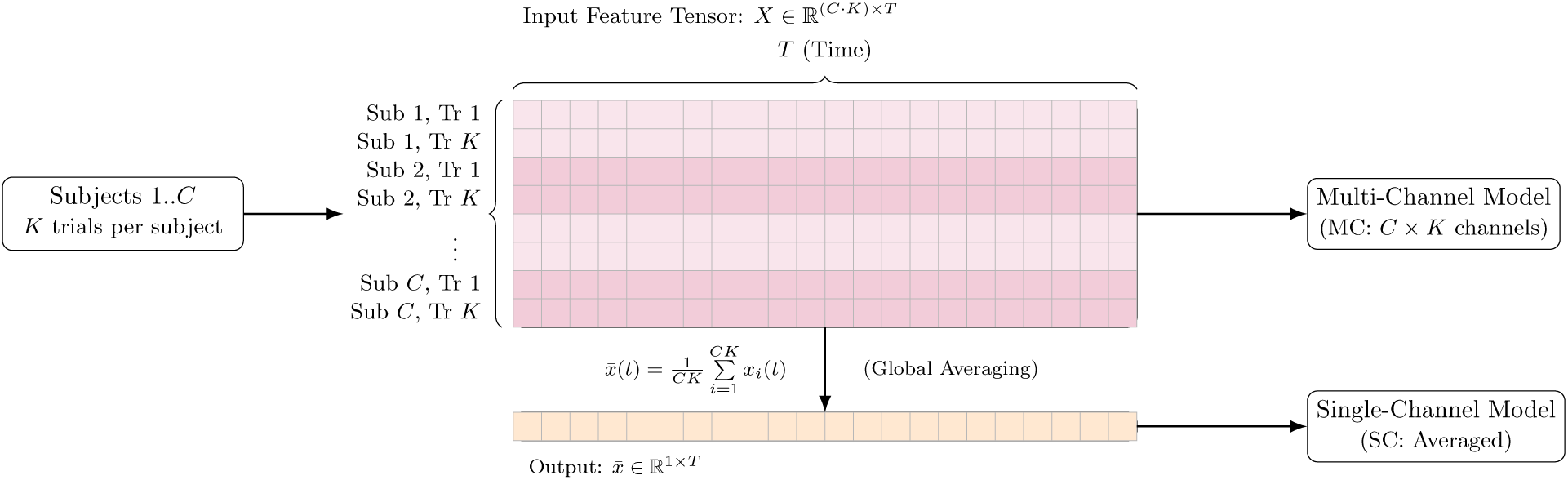
Combined cross-subject and cross-trial aggregation used in the *K*-trials experiment (*C* subjects, *K* trials each).

**Table 10:** Cross-subject averaging with *K* trials per participant. ITR is reported in bits per aggre-gated decision.

| Model | $K$ | Ch. | Approach | Accuracy | ITR | 95% CI |
| --- | --- | --- | --- | --- | --- | --- |
| EEGNet | 2 | 20 | MC | 0.986 | 0.89 | [0.976, 0.996] |
| EEGNet | 2 | 20 | SC | 0.934 | 0.65 | [0.912, 0.956] |
| BaseCNN | 2 | 20 | MC | 0.952 | 0.72 | [0.933, 0.971] |
| BaseCNN | 2 | 20 | SC | 0.930 | 0.63 | [0.907, 0.952] |
| SVM | 2 | 20 | averaging | 0.945 | 0.69 | [0.925, 0.965] |
| EEGNet | 3 | 30 | MC | 0.994 | 0.95 | [0.986, 1.000] |
| EEGNet | 3 | 30 | SC | 0.952 | 0.72 | [0.929, 0.975] |
| BaseCNN | 3 | 30 | MC | 0.967 | 0.79 | [0.948, 0.986] |
| BaseCNN | 3 | 30 | SC | 0.976 | 0.84 | [0.959, 0.992] |
| SVM | 3 | 30 | averaging | 0.982 | 0.87 | [0.967, 0.996] |

The combined approach (Table 10) showed the best results among all tested strategies in the no-aperture condition. At *K* = 3 (30 channels), multi-channel EEGNet achieved an ITR of 0.95 bits/aggregated decision, approaching the theoretical maximum of 1 bit per binary decision. The same strategy further improved on Aperture data (Table 13), where EEGNet reached ITR = 0.97 bits/aggregated decision at *K* = 3. Even at *K* = 2 (20 channels), EEGNet MC showed an ITR of 0.89 bits/aggregated decision, significantly exceeding all previous approaches including cross-subject averaging without repetitions (0.82). SVM with averaging of 30 channels achieved an ITR of 0.87 bits/aggregated decision, demonstrating that even a simple classifier can achieve high quality with sufficient aggregated information.

### 4.9 Time-Shifted Channels

In addition to aggregating multiple stimulus repetitions or multiple subjects, we explored a fundamentally different strategy: constructing multi-channel input from a *single* trial by extracting overlapping time-shifted windows from an extended EEG epoch. This approach does not require additional stimulus repetitions or data from other participants; instead, it exploits the temporal extent of the post-stimulus response within a single recording.

For this experiment, we used the full 2-second (500 samples at 250 Hz) post-stimulus epochs from the same dataset. From each 500-sample epoch, *N* overlapping windows of 250 samples (1 s) were extracted, each shifted by 100 ms (25 samples) relative to the previous one. For *N* = 5:

- Channel 0: samples 0–250 (0–1000 ms)
- Channel 1: samples 25–275 (100–1100 ms)
- Channel 2: samples 50–300 (200–1200 ms)
- Channel 3: samples 75–325 (300–1300 ms)
- Channel 4: samples 100–350 (400–1400 ms)

This yields a sample of shape *X* ∈ R^5×250^, where each channel captures slightly different temporal dynamics of the same stimulus response. As in previous experiments, both MC (raw multi-channel) and SC (averaged) variants were evaluated for neural networks, and SVM was applied to the averaged representation.

The tensor layout for this setting is shown in Fig. 10. Unlike repetition-based aggregation, the channels are highly correlated because they come from overlapping windows of one epoch. This correlation structure is important for interpreting the weaker gains of time-shifted aggregation relative to true repeated-trial averaging.

**Figure 10:**
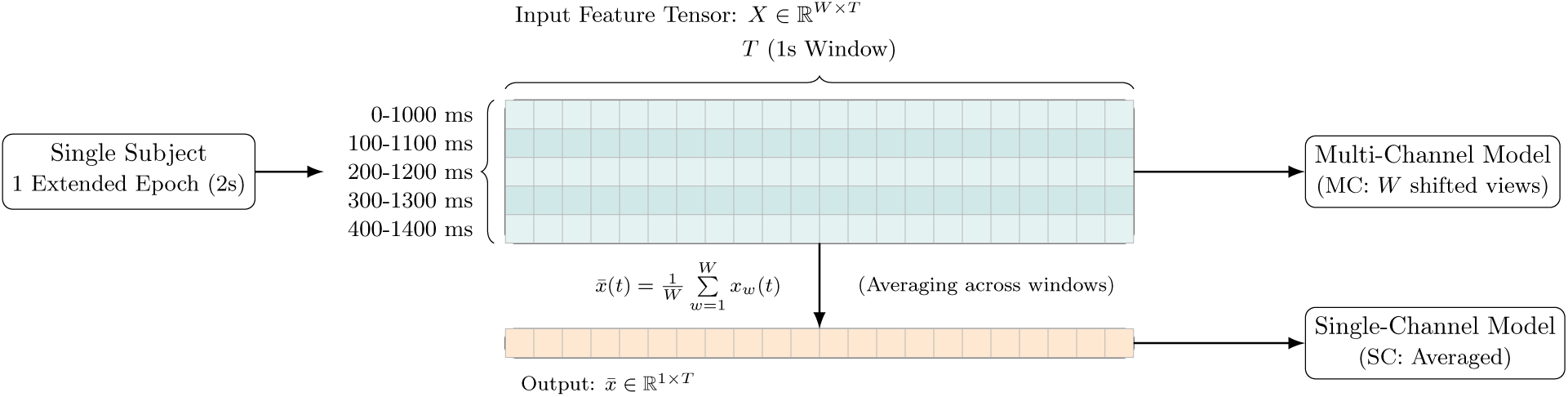
Time-shifted channel construction from one extended epoch: overlapping windows form MC input, or are averaged for SC input.

Results are presented in Tables 11 and 12. The time-shifted MC EEGNet achieved a mean ITR of 0.39 bits/decision unit, comparable to single-trial EEGNet performance (0.35, Table 1). This suggests that the overlapping temporal windows within a single epoch provide only marginally more discriminative information than the original 1-second window. Notably, the SC performance dropped substantially (ITR = 0.17 for EEGNet), indicating that averaging the shifted windows degrades the signal. This is expected: unlike epoch averaging, where multiple independent presentations of the same stimulus reinforce the coherent P300 component, time-shifted windows are highly correlated, and their average dilutes the P300 peak rather than enhancing it.

**Table 11:** Multi-channel (MC) models with *N* = 5 time-shifted windows (shift = 100 ms). SVM uses the averaged representation. ITR is reported in bits per evaluated decision unit.

| Participant | EEGNet |  | BaseCNN |  | SVM |  |
| --- | --- | --- | --- | --- | --- | --- |
|  | Acc. | ITR | Acc. | ITR | Acc. | ITR |
| S0201 | 0.84 | 0.36 | 0.69 | 0.11 | 0.86 | 0.41 |
| S0601 | 0.85 | 0.39 | 0.68 | 0.09 | 0.84 | 0.36 |
| S0701 | 0.81 | 0.30 | 0.65 | 0.07 | 0.84 | 0.36 |
| S1201 | 0.82 | 0.31 | 0.71 | 0.13 | 0.82 | 0.33 |
| S1401 | 0.84 | 0.36 | 0.73 | 0.16 | 0.77 | 0.22 |
| S1601 | 0.86 | 0.43 | 0.78 | 0.24 | 0.83 | 0.34 |
| S1701 | 0.84 | 0.37 | 0.76 | 0.20 | 0.86 | 0.43 |
| S1801 | 0.81 | 0.29 | 0.65 | 0.06 | 0.76 | 0.21 |
| S1901 | 0.97 | 0.82 | 0.92 | 0.60 | 0.93 | 0.63 |
| S2001 | 0.82 | 0.32 | 0.71 | 0.13 | 0.82 | 0.31 |
| **Mean** | **0.85** | **0.39** | **0.73** | **0.18** | **0.83** | **0.36** |

**Table 12:** Single-channel (SC) models with averaging of *N* = 5 time-shifted windows (shift = 100 ms). ITR is reported in bits per evaluated decision unit.

| Participant | EEGNet SC |  | BaseCNN SC |  |
| --- | --- | --- | --- | --- |
|  | Acc. | ITR | Acc. | ITR |
| S0201 | 0.74 | 0.17 | 0.73 | 0.16 |
| S0601 | 0.74 | 0.18 | 0.68 | 0.10 |
| S0701 | 0.72 | 0.14 | 0.66 | 0.08 |
| S1201 | 0.72 | 0.14 | 0.66 | 0.08 |
| S1401 | 0.72 | 0.14 | 0.69 | 0.10 |
| S1601 | 0.77 | 0.23 | 0.71 | 0.13 |
| S1701 | 0.73 | 0.16 | 0.66 | 0.07 |
| S1801 | 0.60 | 0.03 | 0.58 | 0.02 |
| S1901 | 0.86 | 0.42 | 0.83 | 0.35 |
| S2001 | 0.70 | 0.12 | 0.68 | 0.09 |
| **Mean** | **0.73** | **0.17** | **0.69** | **0.12** |

**Table 13:** Aperture data: compact summary for ITR (bits per evaluated decision unit). EEG/Base columns report MC and SC. SVM-I: mean std across subject-specific SVMs (where applicable). SVM-P: pooled SVM on all subjects (or the global SVM where the approach is inherently pooled). For per-subject strategies, EEG/Base report Mean±Std across participants.

| Strategy | EEG-MC | EEG-SC | Base-MC | Base-SC | SVM-I | SVM-P |
| --- | --- | --- | --- | --- | --- | --- |
| Epoch avg., $N = 5$ | 0.540±0.186 | 0.506±0.216 | 0.454±0.222 | 0.438±0.184 | 0.567±0.183 | 0.377 |
| Epoch avg., $N = 10$ | 0.612±0.219 | 0.656±0.221 | 0.568±0.213 | 0.608±0.228 | 0.664±0.243 | 0.505 |
| Cross-subject (10 ch.) | 0.813 | 0.565 | 0.689 | 0.562 | – | 0.572 |
| Mixed, $N = 5$ | 0.318 | 0.399 | 0.352 | 0.287 | – | 0.370 |
| Mixed, $N = 10$ | 0.507 | 0.513 | 0.501 | 0.507 | – | 0.467 |
| $K$ trials × subjects, $K = 2$ (20 ch.) | 0.905 | 0.676 | 0.763 | 0.709 | – | 0.763 |
| $K$ trials × subjects, $K = 3$ (30 ch.) | 0.970 | 0.970 | 0.886 | 0.946 | – | 0.803 |
| Time-shift, $N = 5$ (100 ms) | 0.447±0.147 | 0.206±0.121 | 0.218±0.169 | 0.129±0.132 | 0.332±0.155 | 0.159 |

The common SVM trained on pooled data achieved ITR = 0.20, while individual SVM models averaged ITR = 0.36. The gap between common and individual models is consistent with other experiments, reflecting inter-subject variability.

### 4.10 Aperture data: replication across aggregation strategies

To test whether the observed ordering of aggregation strategies is robust to changes in the recording interface, we repeated the full experimental pipeline on *Aperture data*(Section 2), where a binocular aperture restricts the visual field and reduces the influence of peripheral non-target flashes [23]. Tables 13 and 14 provide a compact summary for ITR and accuracy, respectively. Detailed per-participant metrics and confidence intervals for all approaches are provided in Appendix 6.

**Table 14:** Aperture data: compact summary for accuracy. Notation matches Table 13.

| Strategy | EEG-MC | EEG-SC | Base-MC | Base-SC | SVM-I | SVM-P |
| --- | --- | --- | --- | --- | --- | --- |
| Epoch avg., $N = 5$ | 0.895±0.054 | 0.881±0.068 | 0.860±0.078 | 0.857±0.075 | 0.904±0.053 | 0.845 |
| Epoch avg., $N = 10$ | 0.914±0.063 | 0.926±0.065 | 0.901±0.069 | 0.912±0.069 | 0.926±0.069 | 0.892 |
| Cross-subject (10 ch.) | 0.972 | 0.910 | 0.944 | 0.909 | – | 0.913 |
| Mixed, $N = 5$ | 0.819 | 0.853 | 0.834 | 0.805 | – | 0.842 |
| Mixed, $N = 10$ | 0.892 | 0.894 | 0.890 | 0.892 | – | 0.879 |
| $K$ trials × subjects, $K = 2$ (20 ch.) | 0.988 | 0.941 | 0.961 | 0.949 | – | 0.961 |
| $K$ trials × subjects, $K = 3$ (30 ch.) | 0.997 | 0.997 | 0.985 | 0.994 | – | 0.969 |
| Time-shift, $N = 5$ (100 ms) | 0.866±0.043 | 0.752±0.060 | 0.749±0.090 | 0.691±0.080 | 0.817±0.059 | 0.730 |

## 5 Discussion

The present study addresses the need for comparison of different data aggregation strategies for P300 detection by conducting a controlled evaluation across six experimental paradigms. The results demonstrate a hierarchy of strategies ordered by ITR performance for the evaluated decision units, summarized below.

### Single-trial classification

On single stimulus presentations, SVM with an RBF kernel consistently outperformed both neural network architectures: mean ITR of 0.64 bits/trial (subject-specific) versus 0.35 for EEGNet and 0.15 for BaseCNN. This advantage can be attributed to the effectiveness of kernel methods on low-dimensional, limited-sample data, where deep networks tend to overfit. Generalized models trained on pooled data showed minimal degradation for SVM (ITR = 0.63 vs. 0.64 for individual models), whereas neural networks showed a more pronounced drop, suggesting that SVM features are more robust to inter-subject variability in the single-trial regime.

### Epoch averaging

Multiple class-conditioned stimulus repetitions generally improved performance for all methods. With 10 repetitions and averaging, subject-specific SVM achieved a mean ITR of 0.68 bits/aggregated decision, SC EEGNet achieved 0.64, and SC BaseCNN achieved 0.58. The SC (averaging) approach performed comparably to or better than the MC (multi-channel) approach for within-subject repetitions, indicating that simple averaging effectively preserves the coherent P300 component while canceling uncorrelated noise—consistent with classical ERP theory [21]. This experiment should be interpreted as a controlled estimate of aggregation benefit. In an online BCI, the practical analogue is not averaging known positive and negative labels, but averaging responses associated with repeated presentations of the same candidate stimulus before classification.

### Multi-subject and cross-subject models

The multi-subject model (Section 4.3), where each participant served as a channel, achieved ITR = 0.86 bits/decision unit for EEGNet, demonstrating the potential of structured multi-source aggregation. The cross-subject averaging experiment (Section 4.6) confirmed this finding: MC EEGNet achieved ITR = 0.82. In both cases, the MC approach substantially outperformed SC averaging (ITR = 0.48), indicating that the neural network extracts additional discriminative information from inter-subject differences rather than simply averaging them out. The learned channel weights (Section 4.4) revealed that the model automatically identifies and upweights participants with stronger, more stable P300 responses.

### Mixed vs. controlled averaging

A key finding is that controlled cross-subject averaging (one channel per subject) dramatically outperforms random trial mixing from a common pool. Mixed averaging achieved at best ITR = 0.51 (BaseCNN SC, *N* = 10), compared to 0.82 for cross-subject MC EEGNet (Table 8). The lack of subject structure in the mixed approach means the model cannot learn to weight different sources appropriately, leading to suboptimal signal aggregation. This underscores the importance of preserving the identity of data sources when combining multi-subject EEG data.

### Combined approach

The combined cross-subject approach with *K* trials per participant yielded the highest ITR values in the study. In the no-aperture recordings, EEGNet MC with *K* = 3 (30 channels) achieved ITR = 0.95 bits/aggregated decision (Table 10), and on Aperture data the same setting reached ITR = 0.97 bits/aggregated decision (Table 13), approaching the theoretical maximum for binary classification of the aggregated decision. This confirms a synergistic effect: combining both inter-subject diversity (which captures different aspects of the P300 response) and within-subject repetitions (which enhance SNR through signal averaging) produces better results than either strategy alone. Notably, even SVM with simple averaging of 30 channels achieved ITR = 0.87, demonstrating that with sufficient data aggregation, a classical classifier can rival deep learning performance in this controlled setting.

### Time-shifted channels

The time-shifted approach (Section 4.9), which constructs multi-channel input from overlapping windows within a single 2-second epoch, achieved ITR = 0.39 for MC EEGNet with *N* = 5 windows at 100 ms shift. This is only marginally above single-trial performance (0.35) and substantially below epoch averaging with 5 repetitions (0.49). The SC variant performed even worse (ITR = 0.17), indicating that averaging correlated time-shifted windows is counterproductive—unlike averaging independent stimulus repetitions, where the P300 signal adds coherently while noise cancels. This result confirms that the benefit of epoch averaging stems from the *statistical independence* of repeated presentations, not merely from having multiple temporal views of the same response.

### Model comparison

For MC models, EEGNet systematically outperformed BaseCNN (ITR = 0.95 vs. 0.79 in the best scenario), owing to its depthwise separable convolution architecture that explicitly models spatial and temporal features separately. For SC models with averaging, the gap between architectures narrowed considerably (0.72 vs. 0.84 for *K* = 3), since averaging removes the multi-channel structure that EEGNet exploits. This finding has practical implications: when epoch averaging is used (as in standard P300 spellers), the choice of architecture is less critical; but when multi-channel information is preserved, EEGNet’s specialized design provides a substantial advantage.

### Repetitions and learned representations

In the classical approach, a large number of stimulus repetitions is typically required to achieve reliable P300 detection. In contrast, convolutional neural networks enable effective decoding with a significantly smaller number of repetitions. Importantly, such models may not necessarily extract the classical P300 component directly, but rather identify other spatiotemporal patterns associated with the evoked neural response. Nevertheless, these patterns are still stimulus-dependent and would not emerge in the absence of the corresponding stimulus.

### Limitations

Several limitations should be considered when interpreting these results. First, the dataset included only 10 healthy male participants of similar age, so broader population-level generalization remains to be tested. Second, the present analysis used only the Pz channel, which is informative for P300 but does not capture the full spatial information available in the original recording montage. Third, the repetition and cross-subject aggregation tests are controlled offline constructions: class-conditioned positive/negative grouping is not directly available before classification in an online BCI, and using subjects as channels is best interpreted as multi-source aggregation unless a multi-user deployment scenario is explicitly intended. The reported ITR values should therefore be interpreted as classification-level information scores for the evaluated decision units, not as direct online BCI bitrates without additionally accounting for stimulus timing, repetitions, and the number of data sources. Finally, an independent external dataset or a leave-subject-out validation design would be needed before making stronger claims about generalization to unseen users.

## 6 Conclusions

A comparison of data aggregation strategies has been conducted for P300 event detection using EEGNet, BaseCNN, and SVM on a 10-subject EEG dataset. Across both no-aperture and Aperture data, the main conclusion is that aggregation improves P300 classification most reliably when the structure of the aggregated observations is preserved. Single-trial classification remained limited (ITR ≤ 0.64 bits/trial), whereas repeated-trial averaging, controlled cross-subject aggregation, and especially the combined *K*-trials-per-subject setting substantially increased ITR.

The best-performing configuration was multi-channel EEGNet with *K* = 3 trials per subject, which reached ITR = 0.95 bits/aggregated decision in the no-aperture condition and 0.97 bits/aggregated decision on Aperture data for the aggregated binary decision. Controlled cross-subject aggregation outperformed random mixed averaging, showing that source identity carries useful structure. Conversely, time-shifted windows extracted from a single epoch provided only limited benefit, supporting the interpretation that the main advantage of epoch averaging comes from combining independent stimulus-locked responses rather than simply increasing the number of input channels.

From a practical perspective, SVM with epoch averaging remains a strong and simple option for single-user P300 settings, while EEGNet is most advantageous when structured multi-channel information is preserved. At the same time, the strongest repetition and cross-subject results should be interpreted with their decision units in mind: class-conditioned positive/negative aggregation is an offline analysis, and online P300 systems would instead aggregate responses by repeated candidate stimulus or symbol before making a final classification decision.

## Acknowledgements

This work was supported by the The Ministry of Economic Development of the Russian Federation in accordance with the subsidy agreement (agreement identifier 000000C313925P4H0002; grant No 139-15-2025-012).

## Appendix: Aperture data summaries

The compact ITR and accuracy summaries for the *Aperture data* condition are provided in Tables 13 and 14. These tables report the same strategy ordering as the no-aperture experiments while avoiding duplication of large per-participant tables in the main manuscript. The full per-participant outputs were used to compute the reported mean and standard-deviation summaries and can be generated from the analysis notebooks.

## Appendix: Hyperparameter Tuning

The hyperparameter search process followed this sequence: (1) Borderline-SMOTE parameters were evaluated, (2) EEGNet architecture parameters (*F* 1, *D*, *F* 2) were tuned for single-channel (general) models, (3) EEGNet training hyperparameters (learning rate, weight decay, batch size) were optimized for single-channel models, (4) BaseCNN training hyperparameters were tuned for single-channel models, and finally (5) multi-channel model hyperparameters were optimized for both EEGNet and BaseCNN architectures. SMOTE parameters were re-evaluated with the optimal EEGNet architecture configuration (*F* 1 = 8, *D* = 8, *F* 2 = 8) to ensure consistency.

### Borderline-SMOTE Parameters

We conducted a grid search over Borderline-SMOTE hyperparameters *k* (number of minority neighbors for synthesis) and *m* (number of neighbors for danger detection). Table 16 presents the results for EEGNet with architecture parameters *F* 1 = 8, *D* = 8, *F* 2 = 8. We report accuracy, ITR (computed from accuracy), and F1-score (to reflect class imbalance). The selected configuration was *k* = 20, *m* = 40.

**Table 15:** Qualitative ranking of aggregation strategies by ITR performance for the evaluated decision units.

| Rank | Strategy class | Interpretation |
| --- | --- | --- |
| 1 | Combined $K$ trials × subjects | Highest controlled aggregation performance; preserves source identity and increases SNR. |
| 2 | Cross-subject MC aggregation | Strong multi-source performance when subject identity is preserved as channel structure. |
| 3 | Within-subject epoch averaging | Practical single-user strategy; averaging repeated stimulus presentations improves reliability. |
| 4 | Mixed averaging | Moderate gains; uncontrolled source mixing weakens aggregation benefits. |
| 5 | Time-shifted channels | Limited benefit because overlapping windows are strongly correlated. |
| 6 | Single-trial models | Baseline regime with limited ITR under noisy single-stimulus decisions. |

**Table 16:** Borderline-SMOTE hyperparameter tuning results for general EEGNet model.

| $k$ | $m$ | Accuracy | ITR | F1-Score |
| --- | --- | --- | --- | --- |
| 5 | 10 | 0.822 | 0.324 | 0.172 |
| 5 | 20 | 0.813 | 0.305 | 0.166 |
| 5 | 40 | 0.830 | 0.342 | 0.166 |
| 10 | 10 | 0.850 | 0.390 | 0.174 |
| 10 | 20 | 0.847 | 0.383 | 0.165 |
| 10 | 40 | 0.857 | 0.408 | 0.163 |
| 15 | 10 | 0.858 | 0.411 | 0.182 |
| 15 | 20 | 0.852 | 0.395 | 0.170 |
| 15 | 40 | 0.869 | 0.440 | 0.166 |
| 20 | 10 | 0.873 | 0.451 | 0.176 |
| 20 | 20 | 0.862 | 0.421 | 0.164 |
| **20** | **40** | **0.880** | **0.471** | **0.160** |

### EEGNet Architecture Parameters

We performed a grid search over EEGNet architecture hyperparameters *F* 1 (temporal filters), *D* (depth multiplier), and *F* 2 (pointwise filters). Table 17 presents selected results. The optimal configuration was *F* 1 = 8, *D* = 8, *F* 2 = 8.

**Table 17:** Selected EEGNet architecture hyperparameter tuning results. Bold indicates selected configuration.

| $F1$ | $D$ | $F2$ | Accuracy | ITR | F1-Score |
| --- | --- | --- | --- | --- | --- |
| 1 | 1 | 1 | 0.092 | 0.000 | 0.119 |
| 1 | 4 | 2 | 0.603 | 0.031 | 0.143 |
| 2 | 4 | 1 | 0.800 | 0.278 | 0.125 |
| 4 | 8 | 2 | 0.796 | 0.270 | 0.147 |
| 4 | 8 | 4 | 0.746 | 0.182 | 0.157 |
| 8 | 1 | 1 | 0.776 | 0.233 | 0.141 |
| 8 | 4 | 2 | 0.845 | 0.378 | 0.129 |
| 8 | 4 | 4 | 0.807 | 0.292 | 0.153 |
| 8 | 8 | 4 | 0.804 | 0.286 | 0.154 |
| **8** | **8** | **8** | **0.875** | **0.456** | **0.169** |

### Training Hyperparameters

Table 18 presents the results of hyperparameter tuning for learning rate, weight decay, and batch size. The optimal configuration was learning rate 1 × 10^−4^, weight decay 1 × 10^−5^, and batch size 1024.

**Table 18:** Selected training hyperparameter tuning results. Bold indicates selected configuration.

| Learning Rate | Weight Decay | Batch Size | Accuracy | ITR | F1-Score |
| --- | --- | --- | --- | --- | --- |
| $1 \times 10^{-5}$ | $1 \times 10^{-6}$ | 1024 | 0.830 | 0.342 | 0.166 |
| $1 \times 10^{-4}$ | $1 \times 10^{-6}$ | 256 | 0.900 | 0.531 | 0.156 |
| $1 \times 10^{-4}$ | $1 \times 10^{-6}$ | 512 | 0.898 | 0.525 | 0.146 |
| $1 \times 10^{-4}$ | $1 \times 10^{-5}$ | **1024** | **0.876** | **0.459** | **0.169** |
| $1 \times 10^{-3}$ | $1 \times 10^{-6}$ | 256 | 0.907 | 0.554 | 0.148 |
| $1 \times 10^{-3}$ | $1 \times 10^{-6}$ | 512 | 0.928 | 0.627 | 0.103 |

### Multi-Channel EEGNet Hyperparameters

For the multi-channel model, we conducted an extensive hyperparameter search. Table 19 presents selected results. The optimal configuration was *F* 1 = 128, *D* = 1, *F* 2 = 256, achieving accuracy 0.984, ITR 0.882 bits/decision unit, and F1-score 0.814.

**Table 19:** Selected multi-channel EEGNet hyperparameter tuning results. Bold indicates optimal configuration.

| $F1$ | $D$ | $F2$ | Accuracy | ITR | F1-Score |
| --- | --- | --- | --- | --- | --- |
| 4 | 1 | 8 | 0.955 | 0.735 | 0.182 |
| 4 | 2 | 32 | 0.972 | 0.816 | 0.641 |
| 4 | 4 | 32 | 0.975 | 0.831 | 0.706 |
| 8 | 1 | 32 | 0.968 | 0.796 | 0.529 |
| 8 | 4 | 32 | 0.975 | 0.831 | 0.713 |
| 16 | 1 | 64 | 0.978 | 0.847 | 0.750 |
| 16 | 1 | 128 | 0.979 | 0.853 | 0.764 |
| 32 | 1 | 128 | 0.981 | 0.864 | 0.787 |
| 32 | 1 | 512 | 0.982 | 0.870 | 0.795 |
| 64 | 1 | 256 | 0.983 | 0.876 | 0.800 |
| 128 | 1 | 128 | 0.980 | 0.859 | 0.756 |
| **128** | **1** | **256** | **0.984** | **0.882** | **0.814** |
| 128 | 1 | 512 | 0.982 | 0.870 | 0.786 |

